# Deciphering the Caffeine-Specific Neuroprotective Axis: Comparative Docking and Pharmacokinetic Evaluation of the Coffee Phytocomplex

**DOI:** 10.64898/2026.05.05.723029

**Authors:** Eugenio Ragazzi, Giuseppe Zagotto, Giovanni Sartore

## Abstract

**Background:** Epidemiological studies consistently report inverse associations between caffeinated coffee consumption and dementia risk. However, the molecular mechanisms linking coffee-derived phytochemicals to neuroprotection remain only partially understood.

**Objective:** To evaluate, through integrated *in silico* pharmacology, the relative contribution of adenosine receptor modulation versus direct amyloidogenic enzyme and kinase inhibition in mediating the putative neuroprotective effects of major coffee constituents.

**Methods:** Molecular docking analyses were conducted for caffeine, paraxanthine, chlorogenic acid, trigonelline, cafestol, and kahweol against adenosine A2A and A1 receptors (A2AR, A1R), β-secretase 1 (BACE1), glycogen synthase kinase-3β (GSK-3β), and NLRP3 inflammasome components. Docking was performed using the CB-Dock2 platform. Binding affinities, interaction patterns, and ligand efficiency metrics were assessed. Blood–brain barrier permeability and ADMET properties were predicted using pkCSM.

**Results:** Caffeine and paraxanthine demonstrated structurally coherent binding within the orthosteric pockets of A2AR and A1R, supported by favorable predicted blood–brain barrier penetration and high unbound fractions. Ligand efficiency analysis identified adenosine receptors as the most pharmacologically plausible targets for small xanthine derivatives. Although larger phytochemicals exhibited stronger absolute docking scores at BACE1, GSK-3β, and NLRP3, predicted pharmacokinetic constraints suggest a small biological effect due to a limited central exposure.

**Conclusions:** These findings support an adenosine receptor–centered mechanism as the dominant molecular axis linking caffeinated coffee consumption to reduced dementia risk, favoring neuroinflammatory and signaling modulation over direct enzymatic inhibition. Experimental validation is warranted to confirm translational relevance.

**GRAPHICAL ABSTRACT:** 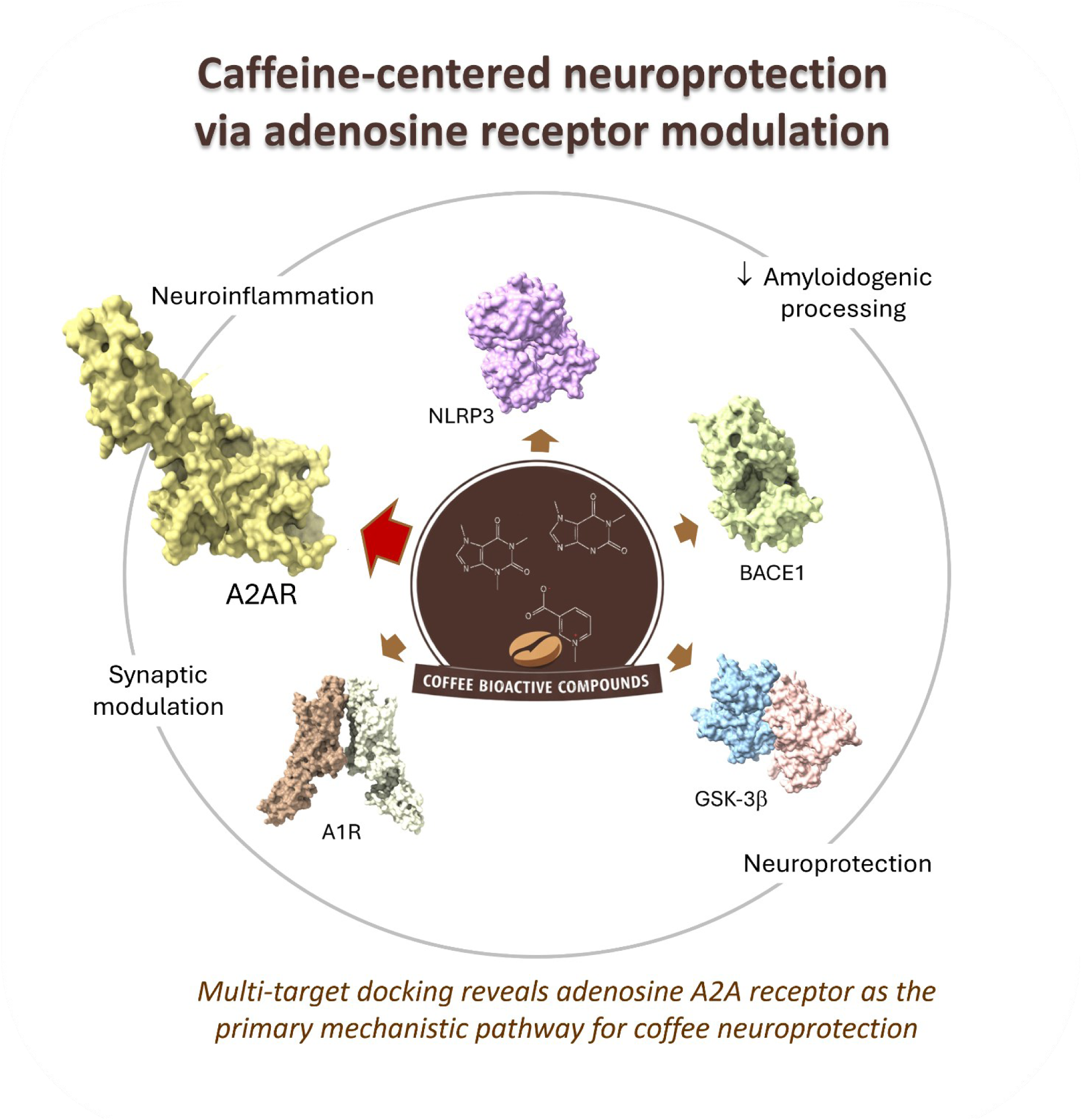

## 1. INTRODUCTION

For years, the association between coffee consumption and reduced cognitive decline was treated as a broad epidemiological observation (Driscoll et al., 2016; Grosso et al., 2017; Kolahdouzan & Hamadeh, 2017; Wang et al., 2024). Notably, prospective data suggest a non-linear relationship between coffee intake and dementia risk, with moderate consumption associated with the greatest protective effect; however, this pattern may reflect a plateau of neuroprotective efficacy at higher intake levels rather than a true increase in risk, as excessive consumption appears primarily to reduce net benefit through systemic adverse effects rather than directly worsening cognitive outcomes (Wang et al., 2024; Zhang et al., 2026). This topic reached a critical turning point with the recent large-scale cohort analyses by Zhang et al. (2026). The observation that neuroprotective associations were confined to caffeinated coffee or tea, and entirely absent in their decaffeinated variants, strongly implicates caffeine as the principal bioactive contributor. This finding shifts the focus from broad lifestyle correlations to a more targeted pharmacological framework. By identifying caffeine as the likely driver of the protective signal, the data underscore the need to delineate its precise molecular mechanisms of action, as well as those of its primary metabolite, paraxanthine. This observation reframes the long-standing “coffee–dementia” association as a pharmacological question rather than a purely nutritional one: namely, whether the epidemiological signal arises from specific adenosine receptor antagonism by methylxanthines rather than from the broader phytochemical matrix of the beverage.

Caffeine (1,3,7-trimethylxanthine), a purine alkaloid of the trimethylxanthine class, is a non-selective antagonist of adenosine receptors, particularly A1 and A2A receptors (Ribeiro & Sebastiao, 2010; Monteiro et al., 2016). A2A receptor (A2AR) plays a central role in regulating synaptic plasticity, microglial activation, dopaminergic signaling, and neuroinflammatory pathways, all processes critically implicated in neurodegeneration (Rivera-Oliver & Díaz-Ríos, 2014; Cunha, 2016; Kolahdouzan & Hamadeh, 2017; Ferré et al., 2018; Ruggiero et al., 2022; Trinh et al., 2022; Salmaso et al., 2025). Also, caffeine intake has been linked to a reduced risk of diabetes (Smith et al., 2006; Biessels, 2010). Genetic inactivation or pharmacological blockade of A2AR confers neuroprotection in Parkinson’s disease models (Chen et al., 2001; Chen & Schwarzschild, 2020), and prospective studies report inverse associations between caffeine intake and Parkinson’s disease risk (Ascherio et al., 2001). Given the shared involvement of adenosinergic dysregulation, neuroinflammation, and synaptic impairment in Alzheimer’s disease (AD), A2AR antagonism (Dall’Igna et al., 2007) has emerged as a plausible mechanistic contributor to dementia risk reduction. However, accumulating evidence indicates that caffeine’s neuroprotective profile may extend beyond A2AR antagonism. Experimental studies show that caffeine reduces amyloid-β (Aβ) accumulation and suppresses β-secretase (BACE1) activity (Arendash et al., 2006, 2009), suggesting that caffeine may interact with multiple molecular targets involved in AD pathogenesis. Additionally, glycogen synthase kinase-3β (GSK-3β), a critical mediator of tau hyperphosphorylation, represents another potential target for multi-mechanistic neuroprotection (Hooper et al., 2008).

Importantly, caffeine is extensively metabolized in the liver via CYP1A2 (Doepker et al., 2016; Szlapinski et al. 2023), with approximately 70-72% converted into paraxanthine (1,7-dimethylxanthine) (Stavric, 1988), the predominant circulating metabolite (Reddy et al., 2024). Because paraxanthine reaches plasma concentrations comparable to caffeine, the “caffeine effect” reported in large-scale cohorts like Zhang et al. (2026) likely reflects a combined parent–metabolite pharmacodynamic profile, as the biological system never encounters caffeine in isolation.

Beyond caffeine, coffee and tea contain a diverse phytocomplex, that is a multi-constituent matrix of bioactive polyphenols, such as chlorogenic acids and flavonoids, that exert known antioxidant and vascular effects, but also neuroprotective properties, including inhibition of amyloid-induced neurotoxicity (Liczbiński & Bukowska, 2022). In pharmacognosy, the therapeutic efficacy of complex plant extracts is often attributed to synergistic interactions between primary active principles and secondary metabolites (Mathur, 2013), suggesting that coffee’s neuroprotective properties could arise from the collective action of its phytocomplex. However, the lack of a neuroprotective association observed with decaffeinated coffee (and tea) still containing the secondary metabolites (Zhang et al., 2026) suggests that within this specific phytocomplex, the non-xanthine components may play a supportive or modulatory role rather than a primary one. This discrepancy necessitates a comparative evaluation to determine whether the caffeine molecule and its metabolites possess a unique pharmacological privileged status across multiple neurodegenerative targets compared to these other major phytochemicals.

Coffee is a chemically complex beverage whose composition depends not only on botanical origin but also on roasting and brewing conditions, which influence the extraction of both volatile and non-volatile constituents (Grosch, 1998; Hu et al., 2019; Cordoba et al., 2020). Major bioactive components include chlorogenic acids, trigonelline, diterpenes such as cafestol and kahweol, and a large number of Maillard-derived products formed during roasting (Hu et al., 2019; Pinheiro et al., 2021). These constituents contribute to the biological profile of coffee and have been associated with antioxidant, anti-inflammatory, and metabolic effects (Socała et al., 2007; Kolb et al., 2020). However, epidemiological evidence showing the absence of neuroprotective association with decaffeinated coffee suggests that, within this complex phytochemical matrix, caffeine and/or its metabolite(s) play the dominant pharmacologically active role in central nervous system effects.

To address this topic, the present study performs a comparative *in silico* pharmacological evaluation of caffeine, its primary metabolite paraxanthine, and selected coffee-derived phytomolecules. Using multi-target docking analysis, we examine binding interactions across key neurodegenerative and neuroinflammatory targets discussed above, including the A2A and A1 receptors, BACE1, GSK-3β, and the NLRP3 inflammasome (Heneka et al., 2018). Considering the caffeine-specific neuroprotective role reported by Zhang et al. (2026), we hypothesize that the caffeine–paraxanthine axis represents the most pharmacologically coherent mechanistic pathway, with adenosine receptor antagonism serving as a central regulatory node within a broader neuroinflammatory network. This approach enables a direct comparison between methylxanthines and non-xanthine coffee constituents to clarify their relative contribution to the observed epidemiological signal. By integrating receptor-level docking, ligand-efficiency analysis, and pharmacokinetic plausibility, this strategy allows evaluation of whether the caffeine–paraxanthine axis constitutes a pharmacologically privileged pathway capable of explaining the population-level association between caffeinated beverage intake and reduced dementia risk.

## 2. MATERIALS AND METHODS

### 2.1. Ligand Selection and Preparation

The criteria adopted to select the coffee-derived phytomolecules were the documented neuroactive, anti-inflammatory, and metabolic properties, as well as their bioavailability. The ligand panel (**Figure 1**) included caffeine, its primary human metabolite paraxanthine (Stavric, 1988), chlorogenic acid and trigonelline as major non-volatile coffee constituents (Cano-Marquina et al., 2013; Heo et al., 2020; Viencz et al., 2023), and the coffee-specific diterpenes cafestol and kahweol, which exhibit reported antioxidative and anti-inflammatory activities (Ren et al., 2019; Lee & Jeong, 2007). Inclusion of paraxanthine enhanced physiological relevance by accounting for caffeine metabolism in humans, while the broader phytochemical panel enabled comparison between xanthine derivatives and structurally distinct coffee compounds. Three-dimensional ligand structures were retrieved from PubChem (https://pubchem.ncbi.nlm.nih.gov/) in SDF format.

**Figure 1.**
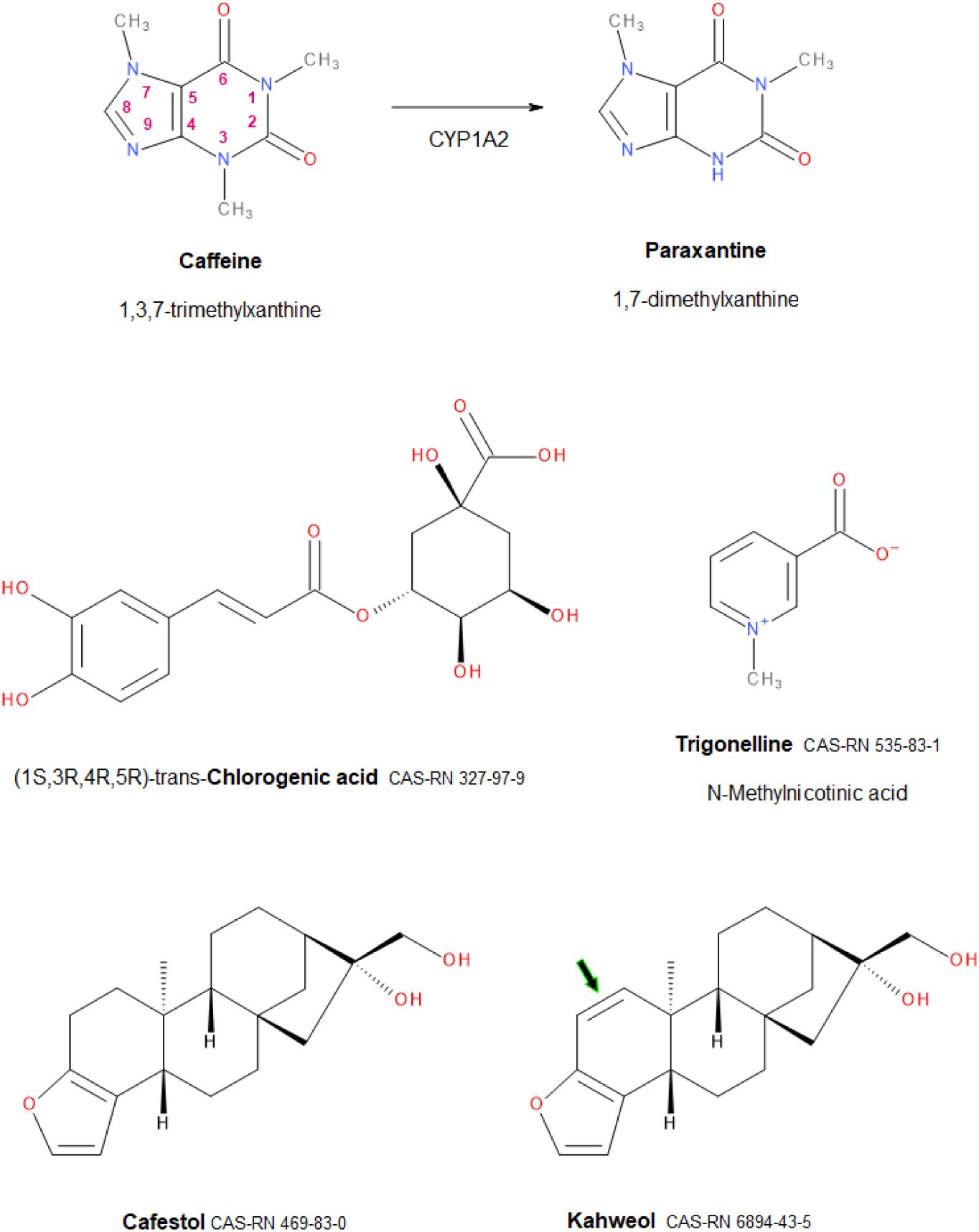
Molecular structures of coffee phytochemicals evaluated for neuroprotective target engagement. Caffeine (1,3,7-trimethylxanthine) undergoes extensive first-pass metabolism via CYP1A2 to generate paraxanthine (1,7-dimethylxanthine), accounting for ∼70% of caffeine clearance in humans. Chlorogenic acid (5-O-caffeoylquinic acid) and trigonelline (N-methylnicotinic acid) represent major non-volatile coffee constituents with reported anti-inflammatory and metabolic activities. Cafestol and kahweol are pentacyclic diterpenes unique to coffee, differing by a C13=C14 double bond (green arrow). Chemical Abstracts Service Registry Numbers (CAS-RN) facilitate unambiguous compound identification.

### 2.2. Target Selection and Structural Preparation

Considering the multifactorial pathophysiology of dementia, a focused multi-target *in silico* approach was implemented to explore potential interactions across complementary pathogenic pathways, ranging from classical adenosinergic antagonism to modern neuroinflammatory and proteopathic hypotheses (Kolahdouzan & Hamadeh, 2017; Scheltens et al., 2021). The 3D structures of the target proteins were retrieved from the RCSB Protein Data Bank (PDB), prioritizing crystal structures of proteins from *Homo sapiens* in antagonist- or inhibitor-bound conformations to ensure well-defined binding pockets suitable for small-molecule docking.

The human Adenosine A2 receptor (A2AR) was selected as the primary mechanistic anchor due to its established involvement in synaptic plasticity and neuroinflammatory regulation (Kolahdouzan & Hamadeh, 2017). The A2AR structure co-crystallized with the high-affinity antagonist ZM241385 (PDB: 5IU4, resolution 1.7 Å; Segala et al., 2016) was chosen as the primary docking template due to its superior resolution and well-defined orthosteric binding pocket. To validate the reproducibility of xanthine binding predictions, A2AR docking was repeated using an alternative crystal structure co-crystallized with caffeine (PDB: 5MZP, resolution 2.1 Å; Cheng et al., 2017). The Adenosine A1 receptor (A1R; PDB ID: 5UEN, resolution 3.2 Å; Glukhova et al., 2017) was included to reflect caffeine’s non-selective antagonism across adenosine receptor subtypes. To address the “amyloid hypothesis” (Hardy & Selkoe, 2002; De Strooper & Karran, 2016), amyloid precursor protein cleaving enzyme 1 (BACE1; PDB ID: 2ZHV, resolution 1.9 Å; Shimizu et al., 2008) was incorporated as a key enzyme in amyloid generation. Furthermore, glycogen synthase kinase-3β (GSK-3β; PDB ID: 1Q5K, resolution 1.9 Å; Bhat et al., 2003) was selected as a critical regulator of tau protein phosphorylation, representing a pivotal node in neurofibrillary degeneration (Hu et al., 2009). Finally, to investigate the role of the innate immune response and microglial activation in dementia progression, the NLRP3 inflammasome (PDB ID: 7ALV, resolution 2.8 Å; Dekker et al., 2021) was included as a representative target for chronic neuroinflammation (McManus & Latz, 2024).

### 2.3. Molecular Docking Procedure

Molecular docking simulations were performed using the web-based platform CB-Dock2 (http://183.56.231.194:8001/cb-dock2/php/index.php), which integrates automated cavity detection with AutoDock Vina scoring (Trott & Olson, 2010). The details of the procedure have been published (Liu et al., 2022). Blind docking was applied to allow unbiased identification of potential binding pockets. For each receptor–ligand pair, the top-ranked cavity based on predicted binding affinity was selected for analysis. Docking conditions were kept identical for all ligands to ensure methodological consistency and comparability. Binding affinity was expressed as Vina score (kcal/mol), and interaction profiles were examined in terms of hydrogen bonding, hydrophobic contacts, cavity volume, and key interacting residues.

Comparative structural analyses evaluated orthosteric alignment of caffeine and paraxanthine within adenosine receptor binding sites and assessed active-site engagement for BACE1, GSK-3β and NLRP3. Docked complexes were visualized using UCSF ChimeraX (Pettersen et al, 2021), and two-dimensional interaction diagrams were generated to characterize hydrogen bonds, π–π stacking interactions, and hydrophobic contacts.

### 2.4. Ligand Efficiency Calculation

To correct for potential scoring bias favoring larger molecules, ligand efficiency (LE) was calculated for the top binding pose as the ratio between the Vina score and the number of non-hydrogen atoms (heavy-atoms, HA), following the definition by Hopkins et al. (Hopkins et al., 2004; Abad-Zapatero, 2007), substituting the experimental ΔG with the binding affinity predicted by AutoDock Vina:

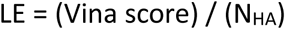

where NHA represents the number of non-hydrogen atoms in the ligand. NHA was obtained for each ligand by PubChem under section “Computed properties”. LE was expressed as kcal/mol per heavy atom (kcal/mol/NHA) as a negative value, where a more negative score indicates higher binding efficiency per atom. This normalized metric enabled direct comparison between smaller xanthines and larger phytochemicals such as chlorogenic acid and diterpenes, providing a size-adjusted evaluation of binding quality.

### 2.5. ADMET Predictions

The pharmacokinetic profiles of the selected ligands, i.e. absorption, distribution, metabolism, excretion, and toxicity (ADMET) properties, were estimated using the pkCSM platform (https://biosig.lab.uq.edu.au/pkcsm/prediction) (Pires et al., 2015). Chemical structures were provided in Canonical SMILES format as input. This predictive framework utilizes graph-based structural signatures and machine learning algorithms to model small-molecule behavior, including membrane permeability, blood–brain barrier (BBB) penetration, and potential metabolic liabilities. Specific emphasis was placed on parameters critical for neuroprotective evaluation, such as Caco-2 permeability, intestinal absorption, and CNS penetration (log PS). The theoretical basis and training datasets for these predictors are detailed by Pires et al. (2015).

## 3. RESULTS

### 3.1. Docking Performance Across Neurodegenerative Targets

Six coffee-derived compounds (caffeine, paraxanthine, trigonelline, chlorogenic acid, cafestol, and kahweol) were docked against five protein targets implicated in neurodegeneration and neuroinflammation: A2A and A1 adenosine receptors (A2AR, A1R), BACE1, GSK-3β, and NLRP3. Binding affinity estimated using AutoDock Vina scores are presented in **Table 1**; LE values calculated to normalize affinity relative to molecular size are shown in **Table 2**. The following discussion should be read in conjunction with these tables and the heatmap in **Figure 2**, which illustrates the global behavior of LE.

**Figure 2.**
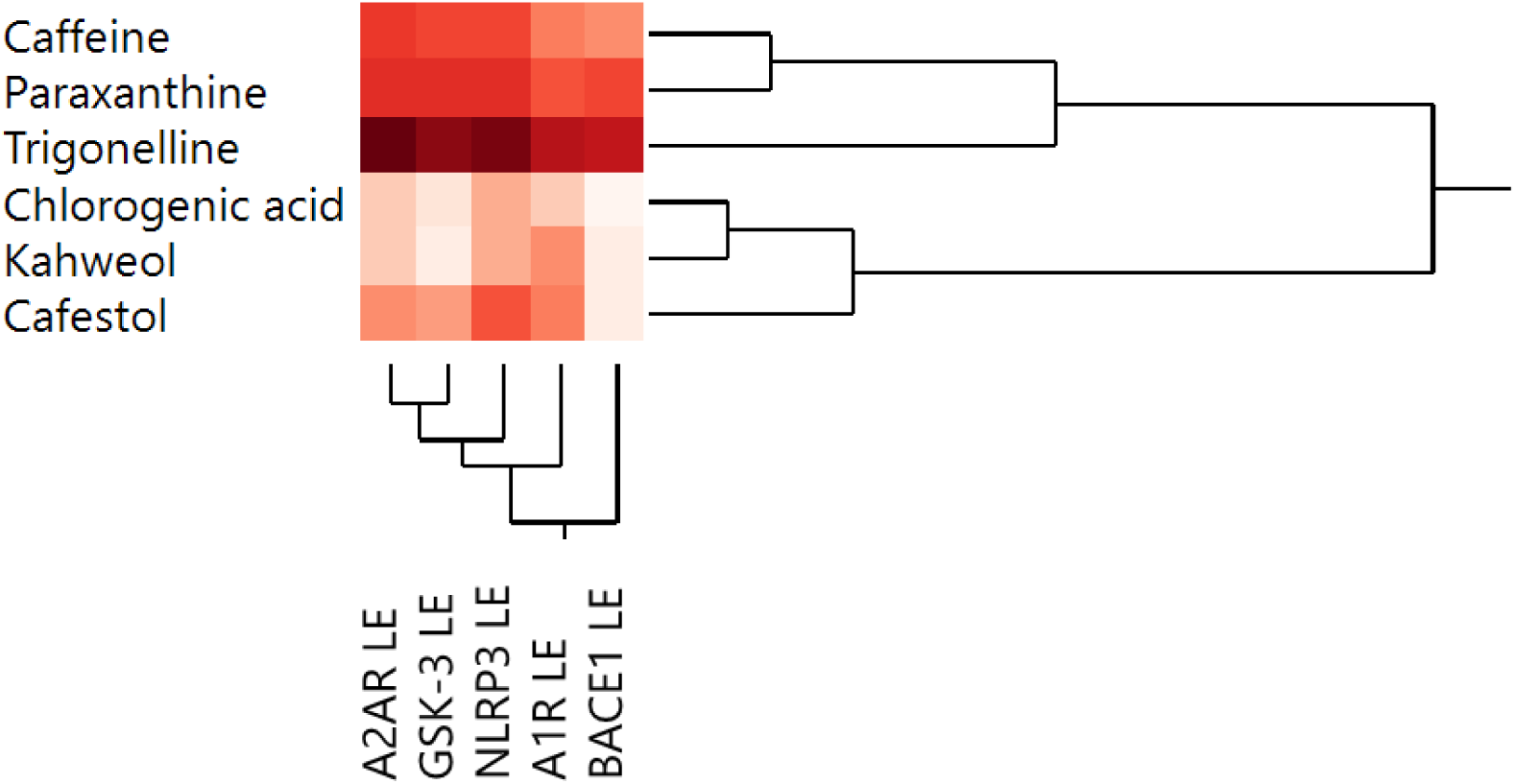
Heatmap and hierarchical clustering (Ward method) of ligand efficiency obtained for the ligands across the selected protein targets. Red color intensity is proportional to the value of ligand efficiency.

**Table 1.**
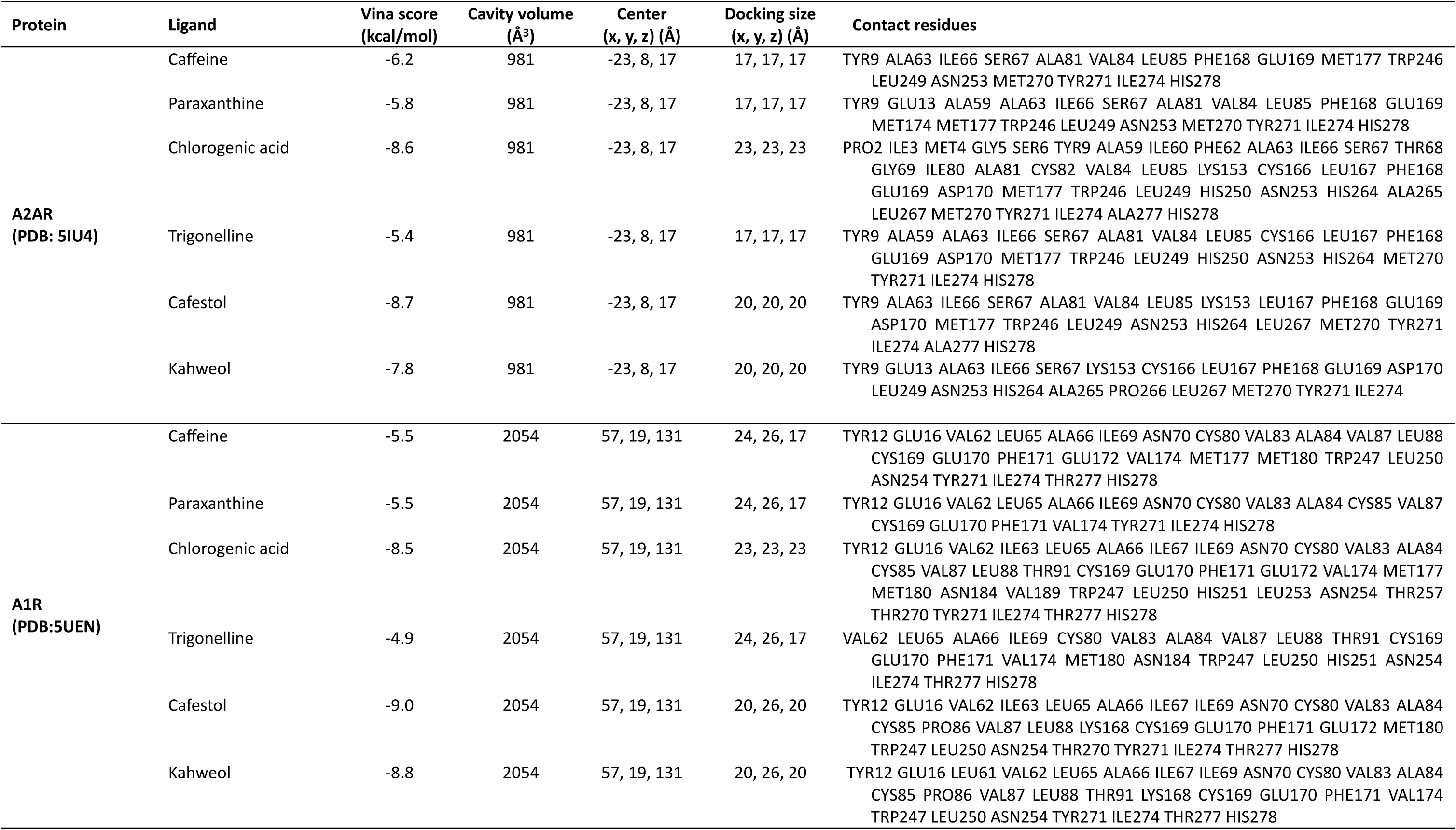
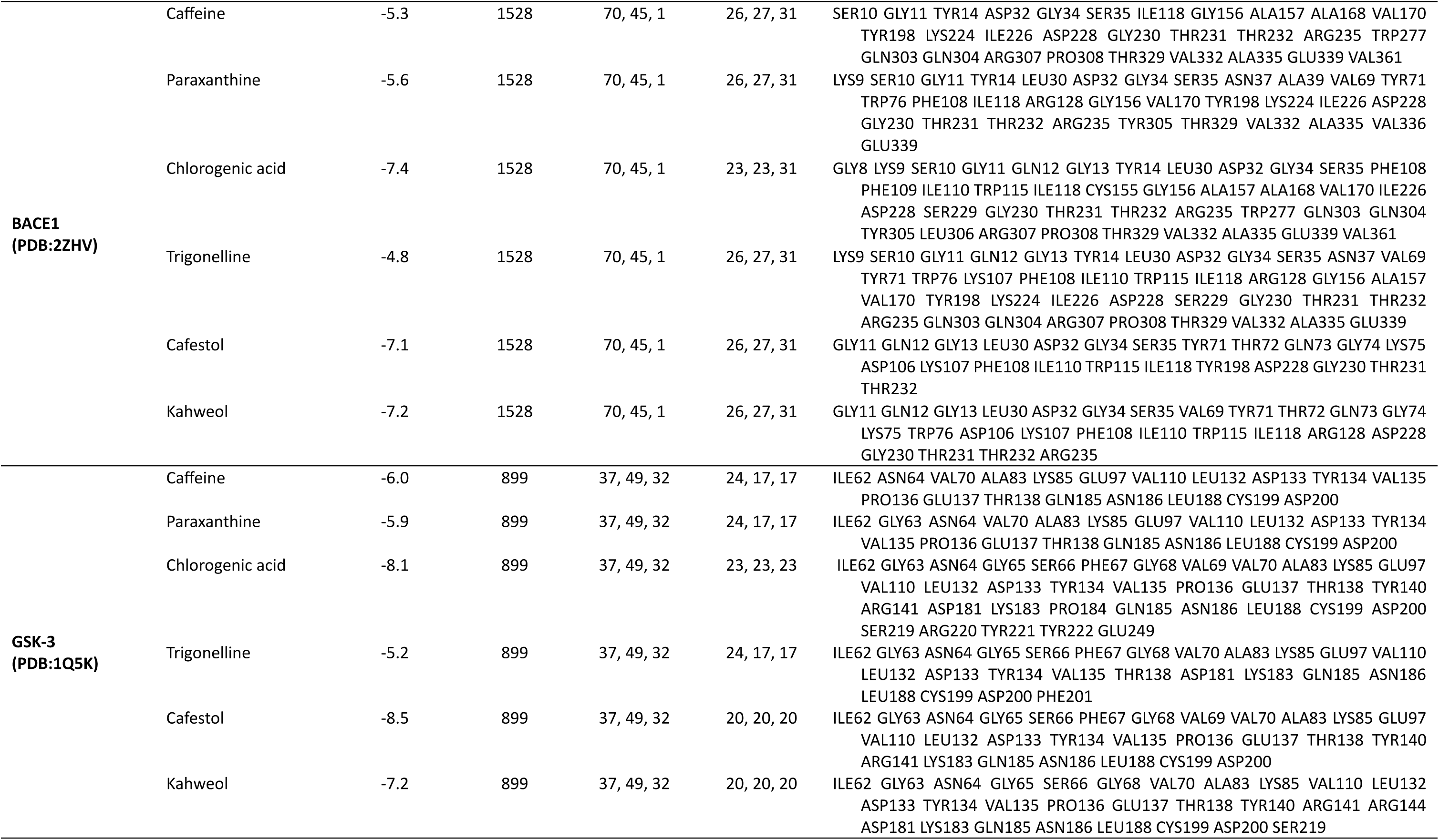
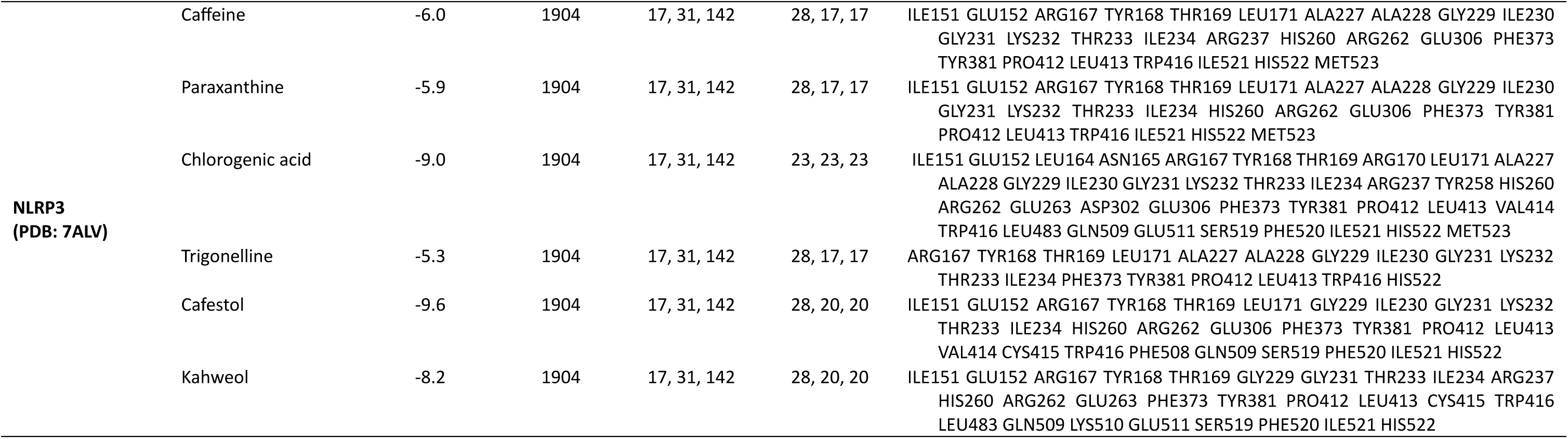
Docking of selected ligands to proteins of interest.

| Protein | Ligand | Vina score (kcal/mol) | Cavity volume (Å<sup>3</sup>) | Center (x, y, z) (Å) | Docking size (x, y, z) (Å) | Contact residues |
| --- | --- | --- | --- | --- | --- | --- |
| **A2AR**<br>**(PDB: 5IU4)** | Caffeine | -6.2 | 981 | -23, 8, 17 | 17, 17, 17 | TYR9 ALA63 ILE66 SER67 ALA81 VAL84 LEU85 PHE168 GLU169 MET177 TRP246 LEU249 ASN253 MET270 TYR271 ILE274 HIS278 |
|  | Paraxanthine | -5.8 | 981 | -23, 8, 17 | 17, 17, 17 | TYR9 GLU13 ALA59 ALA63 ILE66 SER67 ALA81 VAL84 LEU85 PHE168 GLU169 MET174 MET177 TRP246 LEU249 ASN253 MET270 TYR271 ILE274 HIS278 |
|  | Chlorogenic acid | -8.6 | 981 | -23, 8, 17 | 23, 23, 23 | PRO2 ILE3 MET4 GLY5 SER6 TYR9 ALA59 ILE60 PHE62 ALA63 ILE66 SER67 THR68 GLY69 ILE80 ALA81 CYS82 VAL84 LEU85 LYS153 CYS166 LEU167 PHE168 GLU169 ASP170 MET177 TRP246 LEU249 HIS250 ASN253 HIS264 ALA265 LEU267 MET270 TYR271 ILE274 ALA277 HIS278 |
|  | Trigonelline | -5.4 | 981 | -23, 8, 17 | 17, 17, 17 | TYR9 ALA59 ALA63 ILE66 SER67 ALA81 VAL84 LEU85 CYS166 LEU167 PHE168 GLU169 ASP170 MET177 TRP246 LEU249 HIS250 ASN253 HIS264 MET270 TYR271 ILE274 HIS278 |
|  | Cafestol | -8.7 | 981 | -23, 8, 17 | 20, 20, 20 | TYR9 ALA63 ILE66 SER67 ALA81 VAL84 LEU85 LYS153 LEU167 PHE168 GLU169 ASP170 MET177 TRP246 LEU249 ASN253 HIS264 LEU267 MET270 TYR271 ILE274 ALA277 HIS278 |
|  | Kahweol | -7.8 | 981 | -23, 8, 17 | 20, 20, 20 | TYR9 GLU13 ALA63 ILE66 SER67 LYS153 CYS166 LEU167 PHE168 GLU169 ASP170 LEU249 ASN253 HIS264 ALA265 PRO266 LEU267 MET270 TYR271 ILE274 |
| **A1R**<br>**(PDB: 5UEN)** | Caffeine | -5.5 | 2054 | 57, 19, 131 | 24, 26, 17 | TYR12 GLU16 VAL62 LEU65 ALA66 ILE69 ASN70 CYS80 VAL83 ALA84 VAL87 LEU88 CYS169 GLU170 PHE171 GLU172 VAL174 MET177 MET180 TRP247 LEU250 ASN254 TYR271 ILE274 THR277 HIS278 |
|  | Paraxanthine | -5.5 | 2054 | 57, 19, 131 | 24, 26, 17 | TYR12 GLU16 VAL62 LEU65 ALA66 ILE69 ASN70 CYS80 VAL83 ALA84 CYS85 VAL87 CYS169 GLU170 PHE171 VAL174 TYR271 ILE274 HIS278 |
|  | Chlorogenic acid | -8.5 | 2054 | 57, 19, 131 | 23, 23, 23 | TYR12 GLU16 VAL62 ILE63 LEU65 ALA66 ILE67 ILE69 ASN70 CYS80 VAL83 ALA84 CYS85 VAL87 LEU88 THR91 CYS169 GLU170 PHE171 GLU172 VAL174 MET177 MET180 ASN184 VAL189 TRP247 LEU250 HIS251 LEU253 ASN254 THR257 THR270 TYR271 ILE274 THR277 HIS278 |
|  | Trigonelline | -4.9 | 2054 | 57, 19, 131 | 24, 26, 17 | VAL62 LEU65 ALA66 ILE69 CYS80 VAL83 ALA84 VAL87 LEU88 THR91 CYS169 GLU170 PHE171 VAL174 MET180 ASN184 TRP247 LEU250 HIS251 ASN254 ILE274 THR277 HIS278 |
|  | Cafestol | -9.0 | 2054 | 57, 19, 131 | 20, 26, 20 | TYR12 GLU16 VAL62 ILE63 LEU65 ALA66 ILE67 ILE69 ASN70 CYS80 VAL83 ALA84 CYS85 PRO86 VAL87 LEU88 LYS168 CYS169 GLU170 PHE171 GLU172 MET180 TRP247 LEU250 ASN254 THR270 TYR271 ILE274 THR277 HIS278 |
|  | Kahweol | -8.8 | 2054 | 57, 19, 131 | 20, 26, 20 | TYR12 GLU16 LEU61 VAL62 LEU65 ALA66 ILE67 ILE69 ASN70 CYS80 VAL83 ALA84 CYS85 PRO86 VAL87 LEU88 THR91 LYS168 CYS169 GLU170 PHE171 VAL174 TRP247 LEU250 ASN254 TYR271 ILE274 THR277 HIS278 |

| Protein | Ligand | Vina score (kcal/mol) | Cavity volume (Å³) | Center (x, y, z) (Å) | Docking size (x, y, z) (Å) | Contact residues |
| --- | --- | --- | --- | --- | --- | --- |
| **BACE1**<br>**(PDB:2ZHV)** | Caffeine | -5.3 | 1528 | 70, 45, 1 | 26, 27, 31 | SER10 GLY11 TYR14 ASP32 GLY34 SER35 ILE118 GLY156 ALA157 ALA168 VAL170 TYR198 LYS224 ILE226 ASP228 GLY230 THR231 THR232 ARG235 TRP277 GLN303 GLN304 ARG307 PRO308 THR329 VAL332 ALA335 GLU339 VAL361 |
|  | Paraxanthine | -5.6 | 1528 | 70, 45, 1 | 26, 27, 31 | LYS9 SER10 GLY11 TYR14 LEU30 ASP32 GLY34 SER35 ASN37 ALA39 VAL69 TYR71 TRP76 PHE108 ILE118 ARG128 GLY156 VAL170 TYR198 LYS224 ILE226 ASP228 GLY230 THR231 THR232 ARG235 TYR305 THR329 VAL332 ALA335 VAL336 GLU339 |
|  | Chlorogenic acid | -7.4 | 1528 | 70, 45, 1 | 23, 23, 31 | GLY8 LYS9 SER10 GLY11 GLN12 GLY13 TYR14 LEU30 ASP32 GLY34 SER35 PHE108 PHE109 ILE110 TRP115 ILE118 CYS155 GLY156 ALA157 ALA168 VAL170 ILE226 ASP228 SER229 GLY230 THR231 THR232 ARG235 TRP277 GLN303 GLN304 TYR305 LEU306 ARG307 PRO308 THR329 VAL332 ALA335 GLU339 VAL361 |
|  | Trigonelline | -4.8 | 1528 | 70, 45, 1 | 26, 27, 31 | LYS9 SER10 GLY11 GLN12 GLY13 TYR14 LEU30 ASP32 GLY34 SER35 ASN37 VAL69 TYR71 TRP76 LYS107 PHE108 ILE110 TRP115 ILE118 ARG128 GLY156 ALA157 VAL170 TYR198 LYS224 ILE226 ASP228 SER229 GLY230 THR231 THR232 ARG235 GLN303 GLN304 ARG307 PRO308 THR329 VAL332 ALA335 GLU339 |
|  | Cafestol | -7.1 | 1528 | 70, 45, 1 | 26, 27, 31 | GLY11 GLN12 GLY13 LEU30 ASP32 GLY34 SER35 TYR71 THR72 GLN73 GLY74 LYS75 ASP106 LYS107 PHE108 ILE110 TRP115 ILE118 TYR198 ASP228 GLY230 THR231 THR232 |
|  | Kahweol | -7.2 | 1528 | 70, 45, 1 | 26, 27, 31 | GLY11 GLN12 GLY13 LEU30 ASP32 GLY34 SER35 VAL69 TYR71 THR72 GLN73 GLY74 LYS75 TRP76 ASP106 LYS107 PHE108 ILE110 TRP115 ILE118 ARG128 ASP228 GLY230 THR231 THR232 ARG235 |
| **GSK-3**<br>**(PDB:1Q5K)** | Caffeine | -6.0 | 899 | 37, 49, 32 | 24, 17, 17 | ILE62 ASN64 VAL70 ALA83 LYS85 GLU97 VAL110 LEU132 ASP133 TYR134 VAL135 PRO136 GLU137 THR138 GLN185 ASN186 LEU188 CYS199 ASP200 |
|  | Paraxanthine | -5.9 | 899 | 37, 49, 32 | 24, 17, 17 | ILE62 GLY63 ASN64 VAL70 ALA83 LYS85 GLU97 VAL110 LEU132 ASP133 TYR134 VAL135 PRO136 GLU137 THR138 GLN185 ASN186 LEU188 CYS199 ASP200 |
|  | Chlorogenic acid | -8.1 | 899 | 37, 49, 32 | 23, 23, 23 | ILE62 GLY63 ASN64 GLY65 SER66 PHE67 GLY68 VAL69 VAL70 ALA83 LYS85 GLU97 VAL110 LEU132 ASP133 TYR134 VAL135 PRO136 GLU137 THR138 TYR140 ARG141 ASP181 LYS183 PRO184 GLN185 ASN186 LEU188 CYS199 ASP200 SER219 ARG220 TYR221 TYR222 GLU249 |
|  | Trigonelline | -5.2 | 899 | 37, 49, 32 | 24, 17, 17 | ILE62 GLY63 ASN64 GLY65 SER66 PHE67 GLY68 VAL70 ALA83 LYS85 GLU97 VAL110 LEU132 ASP133 TYR134 VAL135 THR138 ASP181 LYS183 GLN185 ASN186 LEU188 CYS199 ASP200 PHE201 |
|  | Cafestol | -8.5 | 899 | 37, 49, 32 | 20, 20, 20 | ILE62 GLY63 ASN64 GLY65 SER66 PHE67 GLY68 VAL69 VAL70 ALA83 LYS85 GLU97 VAL110 LEU132 ASP133 TYR134 VAL135 PRO136 GLU137 THR138 TYR140 ARG141 LYS183 GLN185 ASN186 LEU188 CYS199 ASP200 |
|  | Kahweol | -7.2 | 899 | 37, 49, 32 | 20, 20, 20 | ILE62 GLY63 ASN64 GLY65 SER66 GLY68 VAL70 ALA83 LYS85 VAL110 LEU132 ASP133 TYR134 VAL135 PRO136 GLU137 THR138 TYR140 ARG141 ARG144 ASP181 LYS183 GLN185 ASN186 LEU188 CYS199 ASP200 SER219 |
| **NLRP3**<br>**(PDB: 7ALV)** | Caffeine | -6.0 | 1904 | 17, 31, 142 | 28, 17, 17 | ILE151 GLU152 ARG167 TYR168 THR169 LEU171 ALA227 ALA228 GLY229 ILE230 GLY231 LYS232 THR233 ILE234 ARG237 HIS260 ARG262 GLU306 PHE373 TYR381 PRO412 LEU413 TRP416 ILE521 HIS522 MET523 |
|  | Paraxanthine | -5.9 | 1904 | 17, 31, 142 | 28, 17, 17 | ILE151 GLU152 ARG167 TYR168 THR169 LEU171 ALA227 ALA228 GLY229 ILE230 GLY231 LYS232 THR233 ILE234 HIS260 ARG262 GLU306 PHE373 TYR381 PRO412 LEU413 TRP416 ILE521 HIS522 MET523 |
|  | Chlorogenic acid | -9.0 | 1904 | 17, 31, 142 | 23, 23, 23 | ILE151 GLU152 LEU164 ASN165 ARG167 TYR168 THR169 ARG170 LEU171 ALA227 ALA228 GLY229 ILE230 GLY231 LYS232 THR233 ILE234 ARG237 TYR258 HIS260 ARG262 GLU263 ASP302 GLU306 PHE373 TYR381 PRO412 LEU413 VAL414 TRP416 LEU483 GLN509 GLU511 SER519 PHE520 ILE521 HIS522 MET523 |
|  | Trigonelline | -5.3 | 1904 | 17, 31, 142 | 28, 17, 17 | ARG167 TYR168 THR169 LEU171 ALA227 ALA228 GLY229 ILE230 GLY231 LYS232 THR233 ILE234 PHE373 TYR381 PRO412 LEU413 TRP416 HIS522 |
|  | Cafestol | -9.6 | 1904 | 17, 31, 142 | 28, 20, 20 | ILE151 GLU152 ARG167 TYR168 THR169 LEU171 GLY229 ILE230 GLY231 LYS232 THR233 ILE234 HIS260 ARG262 GLU306 PHE373 TYR381 PRO412 LEU413 VAL414 CYS415 TRP416 PHE508 GLN509 SER519 PHE520 ILE521 HIS522 |
|  | Kahweol | -8.2 | 1904 | 17, 31, 142 | 28, 20, 20 | ILE151 GLU152 ARG167 TYR168 THR169 GLY229 GLY231 THR233 ILE234 ARG237 HIS260 ARG262 GLU263 PHE373 TYR381 PRO412 LEU413 CYS415 TRP416 LEU483 GLN509 LYS510 GLU511 SER519 PHE520 ILE521 HIS522 |

**Table 2.** Docking scores and ligand efficiency across selected targets.

| Ligand | PubChem CID | Molecular weight | Log P<sup>†</sup> | NHA | A2AR (Vina / LE) | A1R (Vina / LE) | BACE1 (Vina / LE) | GSK-3β (Vina / LE) | NLRP3 (Vina / LE) |
| --- | --- | --- | --- | --- | --- | --- | --- | --- | --- |
| Caffeine | 2519 | 194.19 | -1.0293 | 14 | -6.2 / -0.44 | -5.5 / -0.39 | -5.3 / -0.38 | -6.0 / -0.43 | -6.0 / -0.43 |
| Paraxanthine | 4687 | 180.16 | -1.0397 | 13 | -5.8 / -0.45 | -5.5 / -0.42 | -5.6 / -0.43 | -5.9 / -0.45 | -5.9 / -0.45 |
| Chlorogenic acid | 1794427 | 354.31 | -0.6459 | 25 | -8.6 / -0.34 | -8.5 / -0.34 | -7.4 / -0.30 | -8.1 / -0.32 | -9.0 / -0.36 |
| Trigonelline | 5570 | 137.14 | -1.1254 | 10 | -5.4 / -0.54 | -4.9 / -0.49 | -4.8 / -0.48 | -5.2 / -0.52 | -5.3 / -0.53 |
| Cafestol | 108052 | 316.40 | 3.6393 | 23 | -8.7 / -0.38 | -9.0 / -0.39 | -7.1 / -0.31 | -8.5 / -0.37 | -9.6 / -0.42 |
| Kahweol | 114778 | 314.40 | 3.7199 | 23 | -7.8 / -0.34 | -8.8 / -0.38 | -7.2 / -0.31 | -7.2 / -0.31 | -8.2 / -0.36 |

<sup>†</sup> LogP obtained with pkCSM model prediction.
NHA: Number of heavy atoms; LE: Ligand Efficiency (kcal/mol/NHA); Vina: Vina score obtained with CB Dock2.

Across all targets, high-molecular-weight phytochemicals such as chlorogenic acid, cafestol and kahweol yielded the most potent absolute binding energies; however, these scores were largely driven by their greater number of heavy atoms. In contrast, caffeine and paraxanthine exhibited significantly higher LE, indicating a more optimized fit within the binding pockets relative to their size. **Figure 3** illustrates details of the interaction model obtained for A2AR with caffeine and paraxanthine as the ligands of major specific interest.

**Figure 3.**
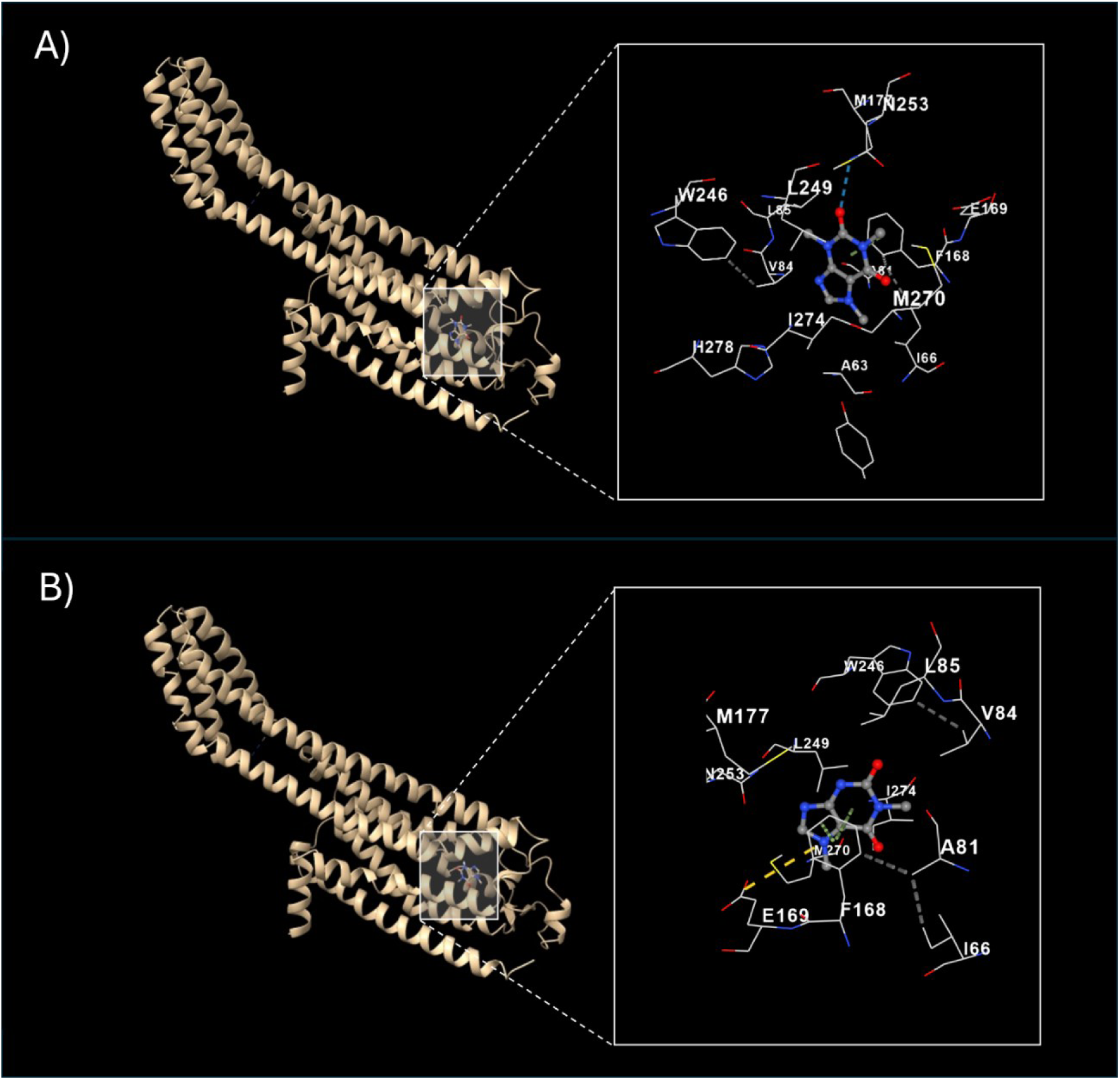
Interaction model obtained for A2AR (PDB: 5IU4) with caffeine (A) and paraxanthine (B). The figure shows cartoon protein-ball-stick interaction model (left), with a detail (right) of the ligand-protein interaction.

### 3.2. Adenosine Receptors (A2AR and A1R)

At A2AR (PDB: 5IU4), cafestol (−8.7 kcal/mol) and chlorogenic acid (−8.6 kcal/mol) displayed the strongest predicted binding, followed by kahweol (−7.8 kcal/mol). Caffeine and paraxanthine showed moderate affinities (−6.2 and −5.8 kcal/mol, respectively), while trigonelline exhibited −5.4 kcal/mol. Docking reproducibility was assessed by comparing predictions against the caffeine-bound A2AR structure (5MZP: **Supplementary Table S1**). Caffeine and paraxanthine binding scores differed by only 0.1 kcal/mol between structures (**Supplementary Table S2**), validating the predicted orthosteric binding mode. Larger phytochemicals exhibited greater score variability (0.6–0.9 kcal/mol), consistent with conformational dependence and reduced pocket complementarity.

At A1R, cafestol (−9.0 kcal/mol), kahweol (−8.8 kcal/mol), and chlorogenic acid (−8.5 kcal/mol) again showed the highest absolute affinities. Caffeine and paraxanthine both yielded −5.5 kcal/mol.

Ligand efficiency analysis shifted the ranking order. Trigonelline demonstrated the highest LE at both A2AR (−0.54) and A1R (−0.49), followed by paraxanthine and caffeine. Larger phytochemicals showed lower efficiency values (approximately −0.39 to −0.34).

Structural inspection of the A2AR docked complex confirmed that caffeine occupies the canonical orthosteric pocket and forms a predicted hydrogen bond with Asn253, together with hydrophobic contacts involving Phe168, Ile274, and Leu249 (**Figure 3**, and **Table 1**). Validation docking using the caffeine-bound structure (5MZP) reproduced these key interactions, with consistent binding pose and contact residues (**Figure S1**), supporting the predicted binding mode.

### 3.3. BACE1, GSK-3β, and NLRP3

Docking to BACE1 revealed stronger absolute affinities for the larger polyphenols, yet paraxanthine achieved the highest LE (−0.43), suggesting a highly specific interaction with the secretase’s catalytic site despite its smaller molecular footprint. Experimental evidence supports xanthine-mediated BACE1 modulation: Janitschke et al. (2019) demonstrated that caffeine and methylxanthines downregulated BACE1 expression and directly decreased β-secretase activity in SH-SY5Y cells, while Sharma et al. (2022) reported that caffeine-based triazoles exhibited dual inhibition of AChE (IC50 1.43 μM) and BACE1 (IC50 10.9 μM).

Similarly, docking to GSK-3β showed the strongest predicted binding for cafestol (−8.5 kcal/mol) and chlorogenic acid (−8.1 kcal/mol), while LE values favored trigonelline (−0.52) and paraxanthine (−0.45). Although caffeine has been proposed to influence neurodegeneration through multiple pathways (Rosso et al., 2008), direct experimental evidence for xanthine-mediated GSK-3β inhibition remains limited, warranting further investigation.

The strongest raw docking scores across the entire study were observed at the NLRP3 inflammasome for cafestol (−9.6 kcal/mol) and chlorogenic acid (−9.0 kcal/mol). Caffeine’s anti-inflammatory effects through NLRP3 modulation have been documented: Wang et al. (2022) showed that caffeine inhibited NLRP3 inflammasome activation via autophagy in microglia from experimental autoimmune encephalomyelitis, while Zhao et al. (2019) demonstrated that caffeine suppressed NLRP3 activation by inhibiting MAPK/NF-κB signaling in LPS-induced macrophages. Nevertheless, the moderate but efficient binding of caffeine and paraxanthine (−6.0 to −5.9 kcal/mol) remains significant when considering the pharmacokinetic constraints of the larger molecules.

### 3.4. In Silico ADMET Prediction

Pharmacokinetic properties were predicted using the pkCSM model (**Table 3**). Caffeine, paraxanthine, and trigonelline exhibited high predicted intestinal absorption (≥94%) and favorable Caco-2 permeability. Crucially, while chlorogenic acid demonstrated strong raw affinity for several targets, its poor predicted intestinal absorption (36.4%) and negligible BBB penetration (log BB −1.41) suggest limited central bioavailability. Conversely, the superior CNS penetration parameters of caffeine and paraxanthine reinforce their roles as the primary active drivers in the caffeinated coffee phytocomplex.

**Table 3.** Prediction of pharmacokinetic (PK) properties (ADMET) of the selected ligands according to pkCSM model.

| ADMET predicted characteristics |  |  | Predicted value |  |  |  |  |  |
| --- | --- | --- | --- | --- | --- | --- | --- | --- |
| PK Property | Model Name | Unit of measure | Caffeine | Paraxanthine | Chlorogenic acid | Trigonelline | Cafestol | Kahweol |
| Absorption | Water solubility | (log mol/L) | -2.023 | -2.389 | -2.449 | -1.931 | -4.087 | -4.403 |
| Absorption | Caco2 permeability | (log Papp in 10<sup>-6</sup> cm/s) | 1.115 | 0.46 | -0.84 | 1.124 | 1.318 | 1.425 |
| Absorption | Intestinal absorption (human) | (% Absorbed) | 99.272 | 93.975 | 36.377 | 96.44 | 94.282 | 94.502 |
| Absorption | Skin Permeability | (log Kp) | -3.376 | -2.735 | -2.735 | -2.736 | -2.956 | -3.171 |
| Absorption | P-glycoprotein substrate | (Yes/No) | No | Yes | Yes | Yes | Yes | Yes |
| Absorption | P-glycoprotein I inhibitor | (Yes/No) | No | No | No | No | No | Yes |
| Absorption | P-glycoprotein II inhibitor | (Yes/No) | No | No | No | No | No | No |
| Distribution | VDss (human) | (log L/kg) | -0.595 | -0.242 | 0.581 | -0.758 | 0.444 | 0.018 |
| Distribution | Fraction unbound (human) | (Fu) | 0.651 | 0.745 | 0.658 | 0.857 | 0.044 | 0.086 |
| Distribution | BBB permeability | (log BB) | -0.268 | -0.569 | -1.407 | -0.234 | -0.234 | 0.272 |
| Distribution | CNS permeability | (log PS) | -2.977 | -3.122 | -3.856 | -2.739 | -1.802 | -1.824 |
| Metabolism | CYP2D6 substrate | (Yes/No) | No | No | No | No | No | No |
| Metabolism | CYP3A4 substrate | (Yes/No) | No | No | No | No | Yes | Yes |
| Metabolism | CYP1A2 inhibitor | (Yes/No) | No | No | No | No | Yes | No |
| Metabolism | CYP2C19 inhibitor | (Yes/No) | No | No | No | No | Yes | No |
| Metabolism | CYP2C9 inhibitor | (Yes/No) | No | No | No | No | No | No |
| Metabolism | CYP2D6 inhibitor | (Yes/No) | No | No | No | No | No | No |
| Metabolism | CYP3A4 inhibitor | (Yes/No) | No | No | No | No | No | No |
| Excretion | Total Clearance | (log ml/min/kg) | 0.193 | 0.355 | 0.307 | 0.378 | 0.57 | 0.526 |
| Excretion | Renal OCT2 substrate | (Yes/No) | No | No | No | No | No | No |
| Toxicity | AMES toxicity | (Yes/No) | No | No | No | No | No | No |
| Toxicity | Max. tolerated dose (human) | (log mg/kg/day) | 0.001 | -0.057 | -0.134 | 0.743 | -1.403 | -0.421 |
| Toxicity | hERG I inhibitor | (Yes/No) | No | No | No | No | No | No |
| Toxicity | hERG II inhibitor | (Yes/No) | No | No | No | No | No | No |
| Toxicity | Oral Rat Acute Toxicity (LD50) | (mol/kg) | 2.802 | 2.184 | 1.973 | 1.878 | 2.954 | 2.826 |
| Toxicity | Oral Rat Chronic Toxicity (LOAEL) | (log mg/kg bw/day) | 0.701 | 1.133 | 2.982 | 0.454 | 1.808 | 1.498 |
| Toxicity | Hepatotoxicity | (Yes/No) | Yes | Yes | No | No | No | No |
| Toxicity | Skin Sensitisation | (Yes/No) | No | No | No | No | No | No |
| Toxicity | *T.Pyriformis* toxicity | (log µg/L) | 0.285 | 0.285 | 0.285 | -0.323 | 0.556 | 0.562 |
| Toxicity | Minnow toxicity | (log mM) | 2.34 | 3.082 | 5.741 | 2.536 | -0.159 | 0.39 |

Predicted blood–brain barrier permeability (as log BB) suggested moderate CNS penetration for caffeine (−0.27) and trigonelline (−0.23), while paraxanthine showed slightly lower permeability (−0.57). However, paraxanthine exhibited a high fraction unbound (0.745), potentially enhancing its availability for target interaction once across the barrier. All compounds except caffeine were predicted to be P-glycoprotein substrates. No compound was predicted to exhibit AMES mutagenicity or hERG channel inhibition. Predicted acute and chronic toxicity values were within ranges compatible with dietary exposure.

## 4. DISCUSSION

The present *in silico* analysis supports adenosine receptor modulation as the primary pharmacological axis through which coffee-derived compounds may influence neurodegenerative pathways. Recent reviews have established adenosine signaling as a critical regulator of neuronal dysfunction and neurodegeneration (Cunha, 2016), with A2AR antagonism by caffeine representing a key mechanism underlying central neuroprotective effects (Ferré et al., 2018). Recent evidence further supports A2AR modulation as a therapeutic target for Alzheimer’s disease through regulation of synaptic plasticity and neuroinflammatory pathways (Trinh et al., 2022). Although larger phytochemicals such as cafestol, kahweol, and chlorogenic acid demonstrated stronger absolute docking scores across multiple targets involved in the neurodegenerative process, ligand efficiency analysis consistently revealed that smaller heterocyclic compounds, particularly caffeine, paraxanthine, and trigonelline, engage binding pockets more efficiently on a size-normalized basis. This distinction is pharmacologically relevant, as orthosteric G protein-coupled receptor (GPCR) sites typically favor compact scaffolds capable of forming optimized interactions within spatially constrained cavities (Zhang et al., 2024).

Caffeine and paraxanthine displayed consistent moderate affinity at both A2A and A1 receptors, supporting their well-established non-selective antagonism (Ribeiro & Sebastiao, 2010). Structural inspection confirmed that caffeine occupies the canonical orthosteric pocket of A2AR, forming a hydrogen bond with Asn253 and engaging Phe168 through π–π stacking, interactions characteristic of competitive antagonists (Doré et al., 2011), and as shown in *in vivo* animal models and confirmed by *in silico* analysis (Doré et al., 2011; Zhong et al., 2024). The observation that paraxanthine maintains, and in some targets exceeds, the ligand efficiency of caffeine suggests that the neuroprotective “signal” (effect) is not only preserved but potentially amplified by CYP1A2 metabolism. This fact could be of interest, since it has been suggested that paraxanthine could be even a safer alternative to caffeine in humans (Szlapinski et al., 2023).

The structural pharmacology of the adenosine A2A receptor has been extensively characterized, with high-resolution crystallographic and computational studies delineating the conformational dynamics and binding determinants of xanthine and non-xanthine antagonists (Segala et al., 2016; Kalash et al., 2017; Cheng et al., 2017; De et al., 2020; Lopes et al., 2021). These investigations have clarified orthosteric pocket architecture, ligand-induced conformational transitions, and key residue networks governing receptor antagonism. Within this established structural framework, the present study does not seek to redefine A2AR binding mechanisms. Rather, it adopts a comparative and integrative perspective, evaluating whether caffeine and its primary metabolite paraxanthine may exhibit a pharmacologically coherent interaction pattern across multiple dementia-relevant targets relative to other major coffee phytochemicals. In this context, docking and ligand-efficiency analyses are interpreted as components of a systems-level hypothesis linking epidemiological associations to receptor-level pharmacology. The high consistency of caffeine and paraxanthine docking scores across two A2AR crystal structures, including the experimentally validated caffeine-bound conformation (Cheng et al., 2017; **Supplementary Table S1**), supports the pharmacological relevance of predicted adenosine receptor interactions.

Given the central role of A2AR signaling in synaptic plasticity, microglial activation, and neuroinflammatory regulation, these findings reinforce the hypothesis that adenosine receptor antagonism represents the dominant molecular mechanism linking habitual coffee consumption to reduced dementia risk, as shown by the recent epidemiological investigation by Zhang et al. (2026), which supports previous literature (Driscoll et al., 2016; Kolahdouzan & Hamadeh, 2017; Ruggiero et al., 2022; Merighi et al., 2023).

In contrast, docking to BACE1 yielded comparatively modest affinities for xanthines, while larger phytochemicals showed stronger but less efficient binding. A similar pattern was observed for GSK-3β, where absolute affinities were higher for diterpenes and chlorogenic acid, yet ligand efficiency did not indicate clear structural optimization. These findings suggest that while coffee-derived compounds may modulate amyloidogenic processing *in vitro* (Fukuyama et al., 2018), such effects are unlikely to represent the primary *in vivo* mechanism due to the unfavorable central bioavailability of the most potent binders (e.g., chlorogenic acid).

NLRP3 displayed the strongest raw docking scores overall, particularly for cafestol and chlorogenic acid. Interestingly, recent evidence suggests that A2AR antagonism may indirectly modulate NLRP3 inflammasome activity (Merighi et al., 2022), providing a mechanistic link between adenosinergic signaling and neuroinflammatory control. However, size-normalized analysis again favored smaller scaffolds. This pattern suggests that predicted inflammasome binding may partly reflect surface complementarity associated with larger, lipophilic molecules rather than highly optimized interaction networks. Nonetheless, given the established involvement of NLRP3 in neuroinflammatory signaling, secondary modulatory contributions from coffee phytochemicals cannot be excluded.

Overall, the data support a hierarchical multi-target framework in which efficient adenosine receptor modulation constitutes the central mechanistic pathway pathway (Cunha, 2016; Trinh et al., 2022), while structurally diverse coffee phytochemicals such as polyphenols (chlorogenic acids: Ishida et al., 2020) and diterpenes may provide ancillary interactions with inflammatory and kinase-related targets. This integrated profile aligns with epidemiological observations linking chronic coffee, tea, and caffeinated beverage consumption to reduced cognitive decline and supports further experimental validation of adenosine-centered mechanisms in neuroprotection.

The predicted ADMET profile further supports the preferential role of small alkaloids in mediating central pharmacological effects. Caffeine and paraxanthine combine high intestinal absorption, moderate predicted BBB permeability, and relatively high unbound fractions, supporting efficient systemic and CNS exposure. Trigonelline also demonstrated favorable absorption and free fraction, with predicted BBB penetration comparable to caffeine; notably, its high intestinal absorption (96.4%) despite unfavorable lipophilicity for passive diffusion suggests carrier-mediated uptake, consistent with structural similarity to nicotinic acid and potential transport via monocarboxylate or organic cation transporters. In contrast, chlorogenic acid exhibited limited predicted intestinal absorption and poor BBB permeability, suggesting reduced likelihood of direct central activity. Although cafestol and kahweol showed favorable membrane permeability and BBB predictions, their low water solubility, extensive plasma protein binding, and predicted CYP interactions may limit free CNS exposure and increase metabolic liability. Overall, the pharmacokinetic profile is concordant with docking findings, reinforcing caffeine and paraxanthine as the most pharmacologically plausible mediators of adenosine-centered neuroprotective mechanisms, while larger phytochemicals may contribute secondary or peripheral effects.

This integrated pharmacodynamic–pharmacokinetic profile provides a plausible mechanistic explanation for why neuroprotective associations appear restricted to caffeinated coffee in large-scale cohort analyses such as the one recently conducted by Zhang et al. (2026). Importantly, similar epidemiological associations have also been reported for other caffeinated beverages, including tea and caffeinated soft drinks, whereas decaffeinated coffee and caffeine-free beverages do not exhibit comparable protective signals (Zhang et al., 2026). This observation further supports the hypothesis that caffeine itself, rather than the broader phytochemical matrix unique to coffee, represents the principal bioactive driver of the observed association. Tea contains caffeine together with other methylxanthines such as theophylline, which are also capable of interacting with adenosine receptors, suggesting that related receptor-level mechanisms may contribute to the protective epidemiological signal across different caffeinated beverages. Our data suggest that the lack of effect in decaffeinated coffee is not due to a lack of target affinity among its constituents (as chlorogenic acid binds strongly to NLRP3 and A2AR), but rather a lack of bioavailability. The predicted ADMET profile identifies caffeine and paraxanthine as the only constituents capable of crossing the blood-brain barrier in concentrations sufficient to achieve meaningful receptor occupancy.

Several practical considerations warrant attention when translating *in silico* predictions to epidemiological contexts. Coffee is not a chemically standardized preparation; the concentration of caffeine and other bioactive constituents varies substantially depending on botanical variety (*Arabica* vs. *Robusta*), roasting conditions, particle size, extraction pressure, water composition, and brewing time (Cordoba et al., 2020). As a result, the caffeine content of brewed coffee beverages can vary widely across preparation methods and commercial outlets. For example, McCusker et al. (2003) reported caffeine concentrations ranging from 58 to 259 mg per serving across espresso and brewed coffees, while more recent analyses confirm that *Robusta*-dominant blends and certain extraction methods yield substantially higher caffeine concentrations per serving (Olechno et al., 2021). Such heterogeneity introduces pharmacokinetic variability that may influence receptor occupancy and partially account for the non-linear dose–response relationships observed in prospective cohort studies. In this context, the apparent U-shaped association reported in some cohorts (Wang et al., 2024) may instead reflect a plateau of adenosine receptor-mediated neuroprotective effects, whereby increasing caffeine exposure beyond moderate levels does not proportionally enhance receptor occupancy, but rather reduces net clinical benefit through off-target systemic effects.

Importantly, the bioactive phytochemical profile of coffee extends beyond caffeine. Chlorogenic acids, trigonelline, and diterpenes such as cafestol and kahweol display relative thermal stability during roasting and brewing processes (Grzelczyk et al., 2022), indicating that coffee consumption represents exposure to a complex phytochemical matrix. While our pharmacokinetic predictions suggest that caffeine and paraxanthine are the most plausible central mediators, structurally diverse co-constituents may contribute peripheral or modulatory effects, supporting a multi-component but hierarchically organized neuroprotective model.

Polyphenolic constituents of coffee and tea, including chlorogenic acids and flavonoids such as catechins, have been extensively investigated for their neuroprotective properties. Experimental evidence indicates that these compounds can modulate oxidative stress, neuroinflammation, and amyloidogenic pathways, with chlorogenic acid demonstrating a reduction in Aβ plaque burden in preclinical models (Molino et al., 2016; Yuan et al., 2025). However, the epidemiological observation that neuroprotective associations are absent in decaffeinated coffee, despite the preservation of these polyphenolic components, suggests that their contribution may be secondary rather than primary. In this context, polyphenols may act as modulatory or synergistic agents within the coffee phytocomplex, whereas caffeine and its metabolites appear to represent the key pharmacologically active drivers capable of achieving sufficient central exposure to influence receptor-mediated pathways.

Furthermore, chronic coffee consumption may influence drug pharmacokinetics through modulation of cytochrome P450 enzymes and transporter systems. Belayneh and Molla (2020) documented clinically significant interactions affecting absorption, distribution, metabolism, and excretion of co-administered medications, particularly those metabolized by CYP1A2. Such interactions carry implications for elderly populations, who are both at higher dementia risk and more likely to be receiving polypharmacy. While our ADMET predictions identified CYP1A2 inhibition by cafestol, the clinical relevance of phytochemical-mediated drug interactions in the context of habitual coffee intake requires further pharmacoepidemiological investigation. These predictions are also consistent with extensive pharmacokinetic literature demonstrating efficient systemic absorption and hepatic metabolism of caffeine, primarily via CYP1A2, producing paraxanthine as the major circulating metabolite (Grzegorzewski et al., 2022).

Collectively, the present findings should be interpreted not as structural rediscovery, but as an integrative pharmacological synthesis that aligns molecular interaction patterns, pharmacokinetic plausibility, and epidemiological observations into a coherent mechanistic framework.

## 5. CONCLUSION

The present docking analysis supports a mechanistic framework in which the neuroprotective associations observed with habitual coffee consumption are most plausibly mediated through efficient adenosine receptor modulation, particularly via small heterocyclic compounds such as caffeine and its primary metabolite paraxanthine. While larger phytochemicals demonstrated stronger absolute binding energies across multiple targets involved in the neurodegenerative process, ligand efficiency analysis indicates that adenosine receptors represent the most structurally coherent and pharmacologically plausible interaction axis.

Importantly, *in silico* pharmacokinetic predictions further support this interpretation. Caffeine and paraxanthine combine favorable intestinal absorption, moderate predicted blood–brain barrier permeability, and relatively high unbound fractions, supporting effective central exposure. In contrast, larger phytochemicals, despite strong predicted binding, display limitations related to solubility, plasma protein binding, or metabolic liability, potentially reducing their direct CNS contribution.

These findings do not establish functional inhibition or clinical efficacy but generate a pharmacologically plausible hypothesis that adenosine-centered signaling constitutes the dominant molecular pathway linking coffee intake to reduced neurodegenerative risk. Experimental validation through binding assays, receptor functional studies, inflammasome activity assays, and *in vivo* models will be necessary to determine the translational significance of these *in silico* observations.

## Authors’ contributions

The authors take sole responsibility for all aspects of this article, including conception, literature review, analysis, and manuscript preparation.

## Acknowledgements

The authors have no acknowledgements to declare.

## Use of Artificial Intelligence Tools

The authors acknowledge the use of an AI language tool (Lucrez-IA, based on Claude 4.5 Sonnet by Anthropic, developed by the Digital Learning e Multimedia Office of the University of Padova) for literature validation, reference formatting, language editing, and exploration of mechanistic hypotheses through literature-based analysis during the preparation of this manuscript. All AI-generated content was critically reviewed, fact-checked, and validated by the authors, who take full responsibility for the scientific accuracy, integrity, and conclusions presented in this work.

## Data Availability

All data generated or analyzed during this study are included in this article and its supplementary information files. Additional data are available from the corresponding author upon reasonable request.

## SUPPLEMENTARY MATERIALS

**Supplementary Table S1.** Docking validation using the caffeine-bound A2AR structure (PDB: 5MZP). To assess reproducibility of binding predictions, all ligands were re-docked to the adenosine A2A receptor co-crystallized with caffeine (PDB: 5MZP; Cheng et al., 2017; resolution 3.0 Å). Caffeine and paraxanthine exhibited minimal score differences compared to the ZM241385-bound structure (5IU4, Table 1), validating the predicted xanthine binding mode. Larger diterpenes showed greater variability (Δ = 0.6–0.9 kcal/mol), consistent with reduced orthosteric pocket complementarity. Docking parameters and scoring methodology were identical to those described in the Methods section to ensure direct comparability.

| Ligand | Vina score (kcal/mol) | Cavity volume (Å<sup>3</sup>) | Center (x, y, z) (Å) | Docking size (x, y, z) (Å) | Contact residues |
| --- | --- | --- | --- | --- | --- |
| Caffeine | -6.1 | 822 | -23, 8, 16 | 17, 17, 17 | TYR9 ALA63 ILE66 SER67 ALA81 VAL84 LEU85 LEU167 PHE168 GLU169 MET177 TRP246 LEU249 ASN253 LEU267 MET270 TYR271 ILE274 HIS278 |
| Paraxanthine | -5.7 | 822 | -23, 8, 16 | 17, 17, 17 | TYR9 ALA59 ALA63 ILE66 SER67 ALA81 VAL84 LEU85 PHE168 GLU169 MET174 MET177 TRP246 LEU249 ASN253 MET270 ILE274 HIS278 |
| Chlorogenic acid | -8.7 | 822 | -23, 8, 16 | 23, 23, 23 | TYR9 LEU58 ALA59 ILE60 PRO61 PHE62 ALA63 ILE66 SER67 THR68 GLY69 ILE80 ALA81 VAL84 LEU85 LYS153 CYS166 LEU167 PHE168 GLU169 ASP170 MET174 MET177 TRP246 LEU249 HIS250 ASN253 HIS264 ALA265 PRO266 LEU267 MET270 TYR271 ILE274 ALA277 HIS278 |
| Trigonelline | -5.3 | 822 | -23, 8, 16 | 17, 17, 17 | TYR9 ALA59 PHE62 ALA63 ILE66 ILE80 ALA81 VAL84 LEU85 PHE168 GLU169 MET177 TRP246 LEU249 ASN253 MET270 ILE274 HIS278 |
| Cafestol | -7.8 | 822 | -23, 8, 16 | 20, 20, 20 | TYR9 GLU13 ALA63 ILE64 ILE66 SER67 ALA81 VAL84 LEU85 LYS153 CYS166 LEU167 PHE168 GLU169 ASP170 TRP246 LEU249 HIS264 ALA265 PRO266 LEU267 MET270 TYR271 ILE274 VAL275 ALA277 HIS278 |
| Kahweol | -8.4 | 822 | -23, 8, 16 | 20, 20, 20 | TYR9 ALA63 ILE66 SER67 THR68 GLY69 LYS150 LYS153 GLN157 CYS166 LEU167 PHE168 GLU169 ASP170 HIS264 ALA265 PRO266 LEU267 MET270 TYR271 ILE274 |

**Supplementary Table S2.**
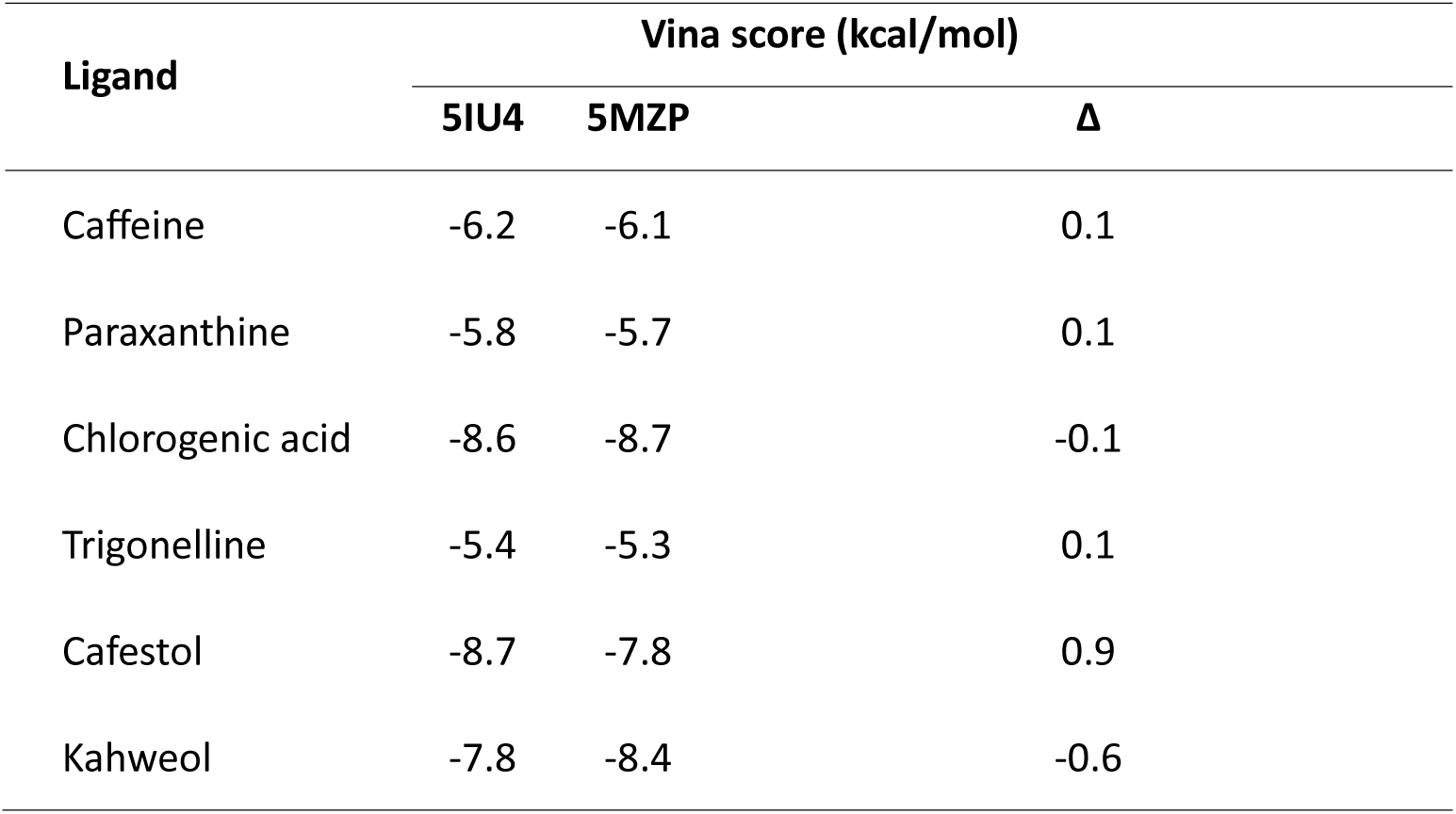
Vina Score Comparison between the two A2AR structures (5IU4 vs 5MZP).

**Supplementary Figure S1.**
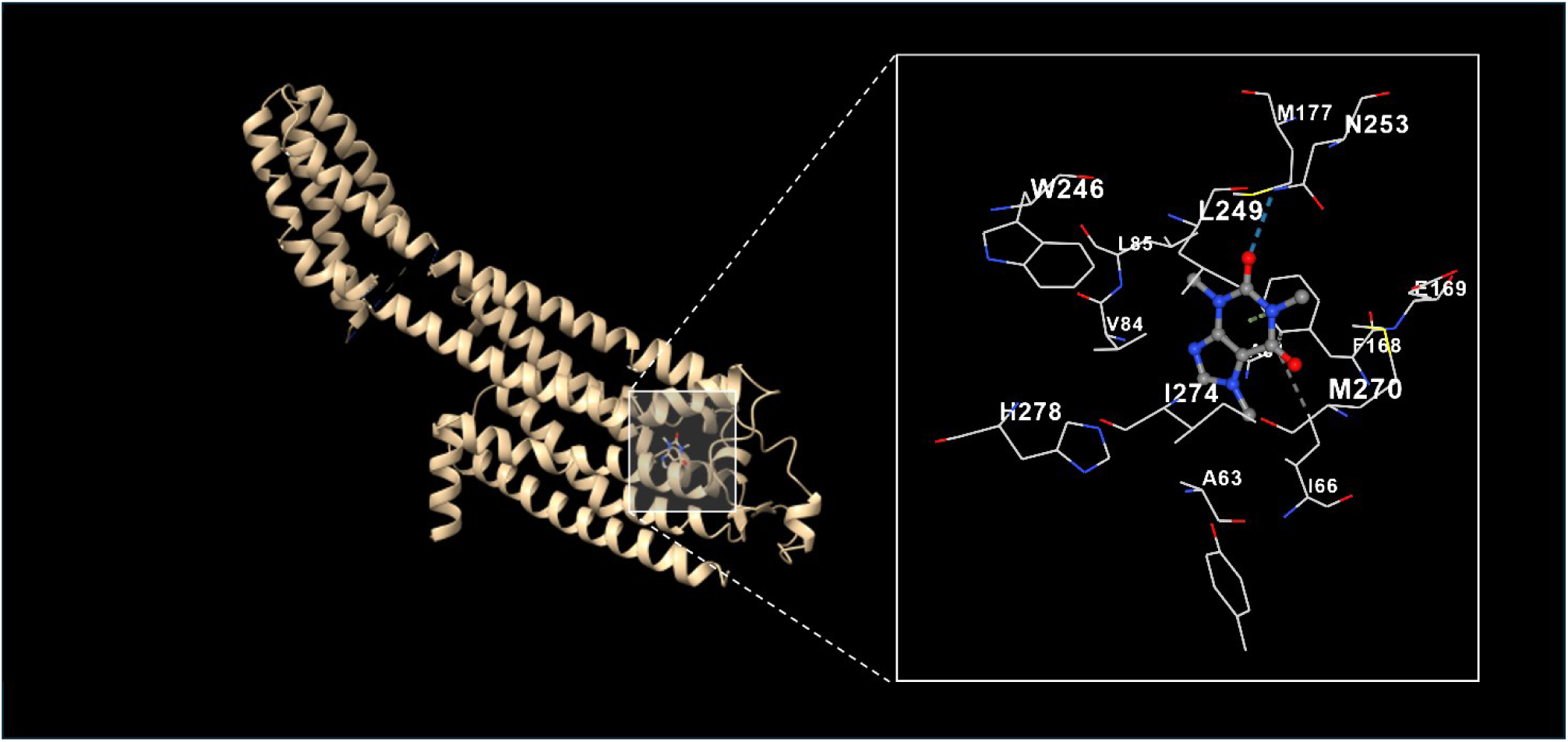
Caffeine binding to A2AR validated with the caffeine-bound crystal structure. Docking of caffeine to A2AR (PDB: 5MZP; Cheng et al., 2017). Left: A2AR transmembrane helices (beige) with caffeine in the binding pocket. Right: Close-up showing caffeine surrounded by key residues including Asn253, Trp246, Phe168, Leu249, and Ile274. The binding pose reproduces key interactions predicted from the antagonist-bound structure (5IU4), confirming binding mode accuracy.

